# In Silico Study of Active Delivery of a Photodynamic Therapy Drug Targeting the Folate Receptor

**DOI:** 10.1101/2025.03.19.644100

**Authors:** Basak Koca Fındık, Elise Lognon, Saron Catak, Antonio Monari

## Abstract

Photodynamic Therapy (PDT), which involves the combined action of a drug and its activation by suitable light, is a particularly attractive novel cancer therapy method due to less systemic side-effects. However, the delivery and accumulation of the PDT drug into cancer cells is still problematic. Here, by using μ-scale molecular dynamic simulations combined with quantum mechanics/molecular mechanics approaches, we examine the behavior of a PDT drug functionalized with a folic acid unit targeting the folate receptor α (FR-α), which is overexpressed in ovarian cancer cells. We show that the PDT drug forms a stable complex with the folate receptor, albeit slightly disrupting the main interaction patterns as compared to the parent folate ligand. Furthermore, we also show that the optical properties of the PDT drug are not altered by its interaction with the protein. Our results confirm that coupling with folate is an attractive strategy for selective active delivery of PDT agents.

## INTRODUCTION

Cancer therapy is a complex and multi-layered problem, usually involving clinical protocols characterized by the combination of diverse treatments, including surgery, chemotherapy, and radiotherapy [1]. Despite significant progresses in recent years, conventional cancer treatments, and especially chemotherapies and radiotherapies, are characterized by heavy secondary effects, which significantly limit the patient’s quality of life and in some cases preclude their use. Chemotherapy or radiotherapy side effects [2] are mainly due to the non-specific action of the drugs or the ionising radiations, which induce cytotoxicity targeting indiscriminately both cancerous and normal cells. Cisplatin [3, 4] developed in the sixties and still in clinical use today can be regarded as a paradigmatic case illustrating the efficacy of chemotherapeutic drugs, but also of the emergence of digestive, respiratory, renal, and even neurological disorders. Furthermore, some chemotherapeutic drugs have also been shown to promote the emergence of cancer cell resistance [5, 6], a situation which can further complicate therapy and lead to cancer relapse.

An appealing alternative to conventional chemotherapy is offered by photodynamic therapy (PDT) in which a photo-sensitive drug is activated by light irradiation in order to exert cytotoxicity on cancer cells [7]. From a photophysical point of view, PDT is based on the excitation of photosensitisers (PS), which can produce reactive oxygen species (ROS), mainly singlet oxygen (^1^O_2_). The high reactivity of ROS will then lead to the disruption of different biological macromolecular structures leading to cell death, ideally by triggering apoptotic pathways. Thus, conventional PDT relies on the triple combination of a suitable PS accumulating in cancer tissues, a light source having a specific wavelength, and the presence of oxygen, which will act as the final cytotoxic agent [8–10]. The main appeal of PDT is that the light-induced drug activation may be precisely controlled spatially, thus significantly reducing general toxicity. Despite its use for the treatment of oral and urogenital cancers, PDT has also shown distinct advantages against bacteria [11–13], fungi [14], viruses [15], and food processing.

Yet, PDT use in clinical practices has remained rather limited [16]. Indeed, the development of an efficient PDT agent is a difficult process requiring to satisfy some different constraints, which may even appear as contradictory [17–19]. As an example, while PDT mostly relies on the activation of ^1^O_2_, yet solid tumours may develop under hypoxic conditions, strongly reducing the efficiency of this approach [20]. Consequently, oxygen-independent approaches based on light activated chemotherapy [21–23], or even molecular photo-switches [24–27] have been recently proposed.

Furthermore, an optimal PS for PDT should absorb in the red and infrared portion of the electromagnetic spectrum, ideally overlapping with the therapeutic window, while favouring photophysical pathways leading to the production of ROS with a high quantum yield. To this end the presence of extended π-conjugated moieties in the PS, such as in porphyrins, may be regarded as ideal. As a matter of fact, hundreds of PS have been proposed and classified as 1^st^ and 2^nd^ generations including porphyrin, chlorin [28, 29] and phthalocyanines [30]. Yet, such compounds may be plagued by aggregation issues, which correlate with low solubility and hence poor bioavailability as well as potential excited state quenching induced by vibrational relaxation. Furthermore, the precise accumulation of the PS at the tumour site is fundamental to maximize its selectivity and therapeutic efficiency. Tackling the problem of bioavailability and reduced solubility is mainly addressed by the so-called 3^rd^ generation PS [8, 9, 31, 32]. Usually this is achieved by a vectorization strategy in which a PS is encapsulated with a drug-delivery agents or is covalently conjugated with a targeting unit [31, 33–35]. Usually hollow molecular structures, such as cyclodextrins, are used as vectorization agents. We have recently shown [29] that the encapsulation of the clinically approved PDT drug Foscan [36] with β-cyclodextrins leads to the formation of a complex, which is rapidly internalized in the cellular membrane where it can spontaneously release the PS. Overall, encapsulation strategy, which is also known as passive targeting, may induce enhanced permeability and retention effects while reducing PS aggregation [37]. Passive delivery is also advantageous, since it protects the PS from degradation, thus increasing its residence time, thereby maximizing treatment efficiency [29, 38, 39]. Yet, passive delivery does not address the necessity to selectively accumulate the drug on tumour sites and cancer cells. A different approach, often referred to as active targeting, relies on the conjugation of the PS with targeting moieties, which are recognized by receptors overexpressed at the surface of cancer cells [40]. Both peptides or small molecules, such as cell penetrating peptide [41], biotin [42], albumin [43], and folic acid [44, 45], have been used to this purpose. Clearly, the receptor-targeted PS should be easily identified and maintain a high binding affinity with the corresponding receptor. Among the targeting moieties, folic acid (FA, Figure 1), which targets folate receptor (FR) is particularly attractive, since FR is overexpressed in various cancer types including lung and ovarian tumours [46, 47].

**Figure 1.**
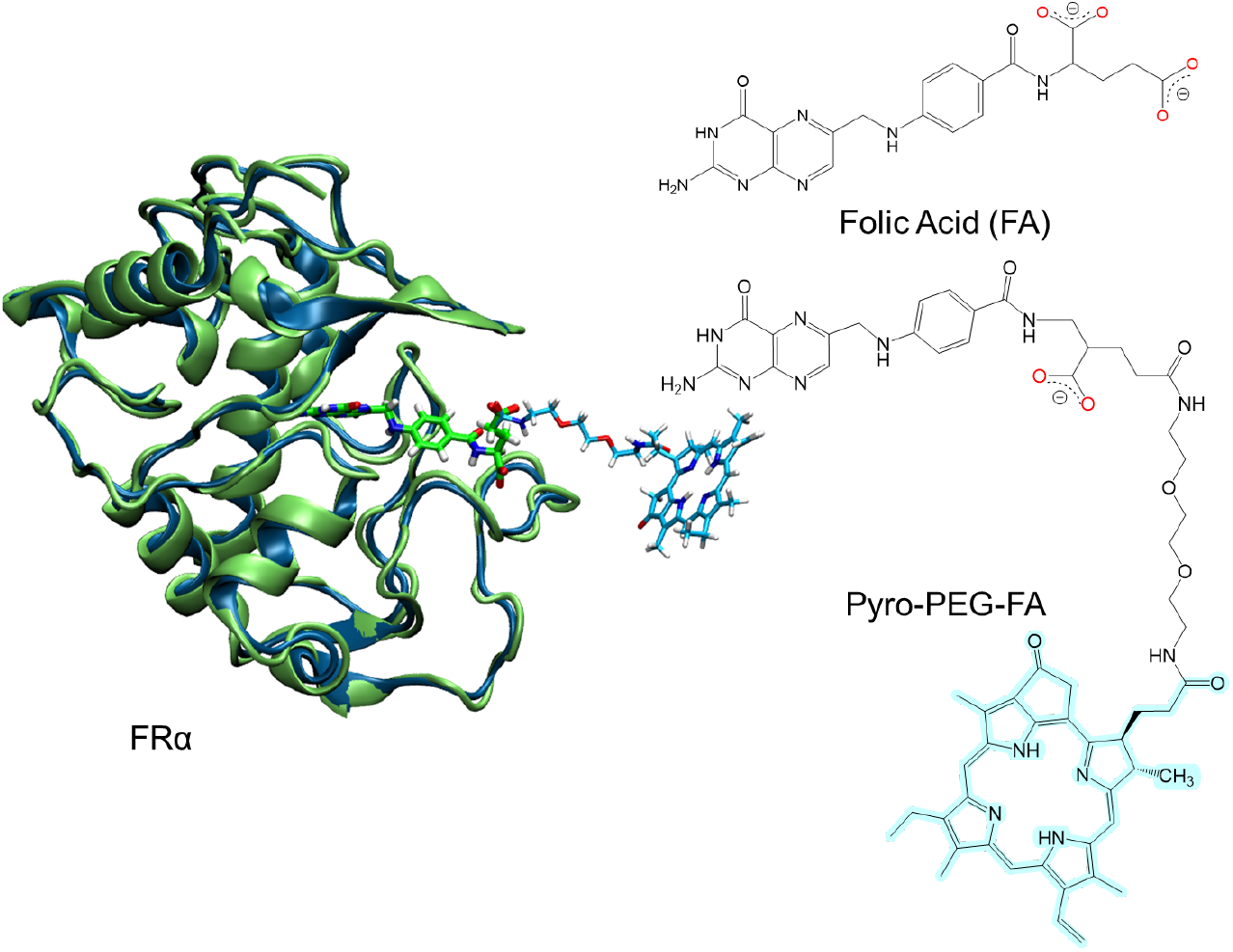
Initial structure of FR bound to FA (green) and Pyro-PEG-FA (blue). The corresponding 2D chemical sketches of the ligands are also given.

FR is a specialized water-soluble protein found at the surface of certain human cells to which it is anchored through a glycosyl phosphatidylinositol (GPI) unit. FR plays a crucial role in the cellular uptake of folate [48], a vital B-vitamin necessary for DNA synthesis, repair, and cell division [49]. Different isoforms of the folate receptor, namely, FR-α, FR-β and FR-γ have been identified [50]. Among them, FR-α is particularly attractive since it is highly expressed in certain cancer cells while is found only in limited quantities on healthy cells [51]. FA fits tightly into the binding pocket of FR-α burying its pterin moiety deep inside the protein cavity while its glutamate-like carboxylic tail interacts with the positively charged FR cavity entrance. Globally, conserved π-stacking and hydrophobic interactions as well as hydrogen bonds contribute to the high affinity and high stability of the FA/FR complex, which presents a dissociation constant K_d_ of about 10^−10^ nM [50].

In 2020 Baydan et al. reported that a porphyrin-based PS (Pyro) could be tethered to FA by a polyethylene glycol linker leading to the Pyro-PEG-FA unit (Figure 1), which could specifically target ovarian tumour cells when irradiated by 672 nm red light [46, 52].

While, the coupling of FA with PS is easy to achieve from a synthetic point of view, the low water solubility of FA and the typically lipophilic nature of tetrapyrrole PS may still limit its bioavailability. While a targeting carrier can enhance the tumour-specific delivery of a photosensitizer (Kd ≈ 0.1∼1 nM), factors such as the carrier’s drug-loading capacity and drug release kinetics still influence its photodynamic effectiveness [44]. Yet the use of a PEG linker to bridge FA with the PS core is ideal to allow the design of a photosensitizer possessing both tumour targeting ability and high-water solubility [53]. The efficiency of the proposed combined Pyro-PEG-FA (Figure 1) was evaluated both *in vivo* and *in vitro* confirming its capacity to significantly induce cancer cell death, and hence reducing the tumour size [46].

To precisely assess the feasibility of the active targeting strategy, it is important to verify that the affinity and selectivity towards the receptors, as well as the inherent spectral and photophysical properties of the tethered PS are preserved. Furthermore, the PS should also be relatively free from the receptor to allow the production of ROS in the vicinity of the cellular membrane. To this effect, the use of molecular modelling and simulation is an appealing approach, since it provides an atomistic-scale resolution of the drug receptor complex in its native configuration as well as in the presence of the tethered PS, while also allowing to compare the binding free energies and assess the properties of the excited state manifold.

In the present study, we aim to clarify the interactions between Pyro-PEG-FA and FR-α, through molecular dynamics (MD) simulations, reaching the μ-second timescale also pointing and compare its behavior with that of the complex between FR-α and the native FA ligand. Furthermore, using hybrid quantum mechanics/molecular mechanics approaches we have also modelled the spectral properties of the chromophoric units of the Pyro-PEG-FA in complex with the receptor and in water solution to assess that the main photophysical and optical properties of the PS are not altered as a consequence of the interaction with the receptor.

## COMPUTATIONAL METHODOLOGY

All-atom MD simulations have been performed via the NAMD 3.0 software [54, 55] to study the structure and properties of FR-α in complex with the native FA as well as the Pyro-PEG-FA drug. The systems are described with standard amber force field parameters, namely ff14SB [56] is used to represent the protein and TIP3P [57] for water. The force fields for the ligands, Pyro-PEG-FA and FA, have been parameterized in the framework of the Generalized AMBER Force Field 2 (GAFF2) [58] procedure for atom type assignments and bond parameters using ANTECHAMBER utility. Atomic point charges have been derived through the restrained electrostatic potential (RESP) protocol [59] based on quantum chemistry calculations of the isolated system to assure accurate charge distributions.

The crystallographic structure of human FR-α in complex with FA (PDB ID: 4LRH [50]) has been used as a starting model. The simulated receptor comprises 204 amino acid residues organized in six α-helices, four β-strands, and several loop regions stabilized by eight disulfide bonds, which have been maintained in the model. Since we are interested in only the extramembrane region of FR the N-acetylglucosamine residues present in the crystal structure close to Asn47, Asn139, and Asn179 were removed. The protonation states of the ionizable amino acids are enforced, according to the dominant species at standard physiological conditions (pH 7.4) as previewed by H++ server [60–62], namely, 20 amino acid residues (13 Glu and 7 Asp) bear a negative charge, and 24 (12 Lys, 11 Arg, and the backbone amine of the N-terminal Gln) are positively charged. To obtain the starting structure of the Pyro-PEG-FA the folate moiety of the latter was aligned to the position of FA in the binding pocket of the crystal structure (Figure 1).

Pyro-PEG-FA/FRα was placed in a 98.0 × 90.0 × 96.0 Å^3^ box and hydrated by 22482 water molecules, while FA/FRα was soaked in a 81.0 × 90.0 × 96.0 Å^3^ water box with 18141 water molecules. In order to adjust the neutral charge state of the overall system, Cl^**-**^ ions have been used.

After the preparation of the initial systems, 60000 conjugated gradient minimization steps were performed before thermalization and equilibration. Temperature and pressure have been set at 300 K and 1 atm and kept constant during the 1µs MD simulation time in a NPT thermodynamic ensemble. Temperature and pressure were controlled by a Langevin thermostat [63] and a Nose-Hoover Langevin barostat [64, 65], respectively. Periodic boundary conditions were used, and electrostatic interactions have been obtained using the Particle Mesh Ewald (PME) [66] summation considering a cut-off of 9 Å. For each simulation, hydrogen mass repartition (HMR) [67] was used in combination with Rattle and Shake algorithm [68] to slow the highest frequency vibrations involving hydrogens, thus allowing the use of a 4.0 fs time step to integrate Newton’s equations of motion. Four different replicas for each system have been performed spanning 1µs each for Pyro-PEG-FA and 500 ns FA. In addition, isolated Pyro-PEG-FA was solvated in a TIP3P water box (79.0 × 56.5 × 53. Å^3^) and its behavior sampled through four independent replicas lasting for 500 ns each in the NPT ensemble.

After performing simulations, Molecular Mechanics Poisson-Boltzmann Surface Area (MMPBSA) and Molecular Mechanics Generalized-Born Surface Area (MMGBSA) were conducted to calculate the enthalpic contribution to the binding free energy calculated for each replica, with the help of MMPBSA.py[69] tool as implemented in Amber22[70].

The absorption spectra of isolated and FR bound Pyro-PEG-FA has been calculated at hybrid quantum mechanics/molecular mechanics (QM/MM) level by obtaining vertical electronic excitations from the ground state *via* time-dependent density functional theory (TD-DFT). To this aim, and using CPPTRAJ tool [71] of AmberTools, 100 snapshots were extracted uniformly along the corresponding trajectories from both environments, and the Terachem/Amber [72] interface was used to perform TD-DFT calculations. Only the pyropheophorbide-a moiety of Pyro-PEG-FA, highlighted in the Figure 1, has been treated at the QM level, using the ωB97XD [73] exchange-correlation functional and the 6-31G(d) basis set, whereas the rest of the molecule, water and FR receptor have been described at classical molecular mechanic levels. The electronic density reorganization in the excited states has been analyzed by means of the Theodore code [74], notably by analyzing Natural Transition Orbitals (NTO) [75].

## RESULTS AND DISCUSSION

Coherently with the observed crystal structure for the FR/FA adduct the binding pocket of FR takes the shape of a long cavity opposing the three α-helices, which is flanked by unstructured loops (Figure 2A). This global pattern is also confirmed in the case of the Pyro-PEG-FA (Figure 2B), which maintains the FA moiety inserted deeply in the binding pocket of the receptor while the PEG arms keeps the Pyro unit further in the bulk.

**Figure 2.**
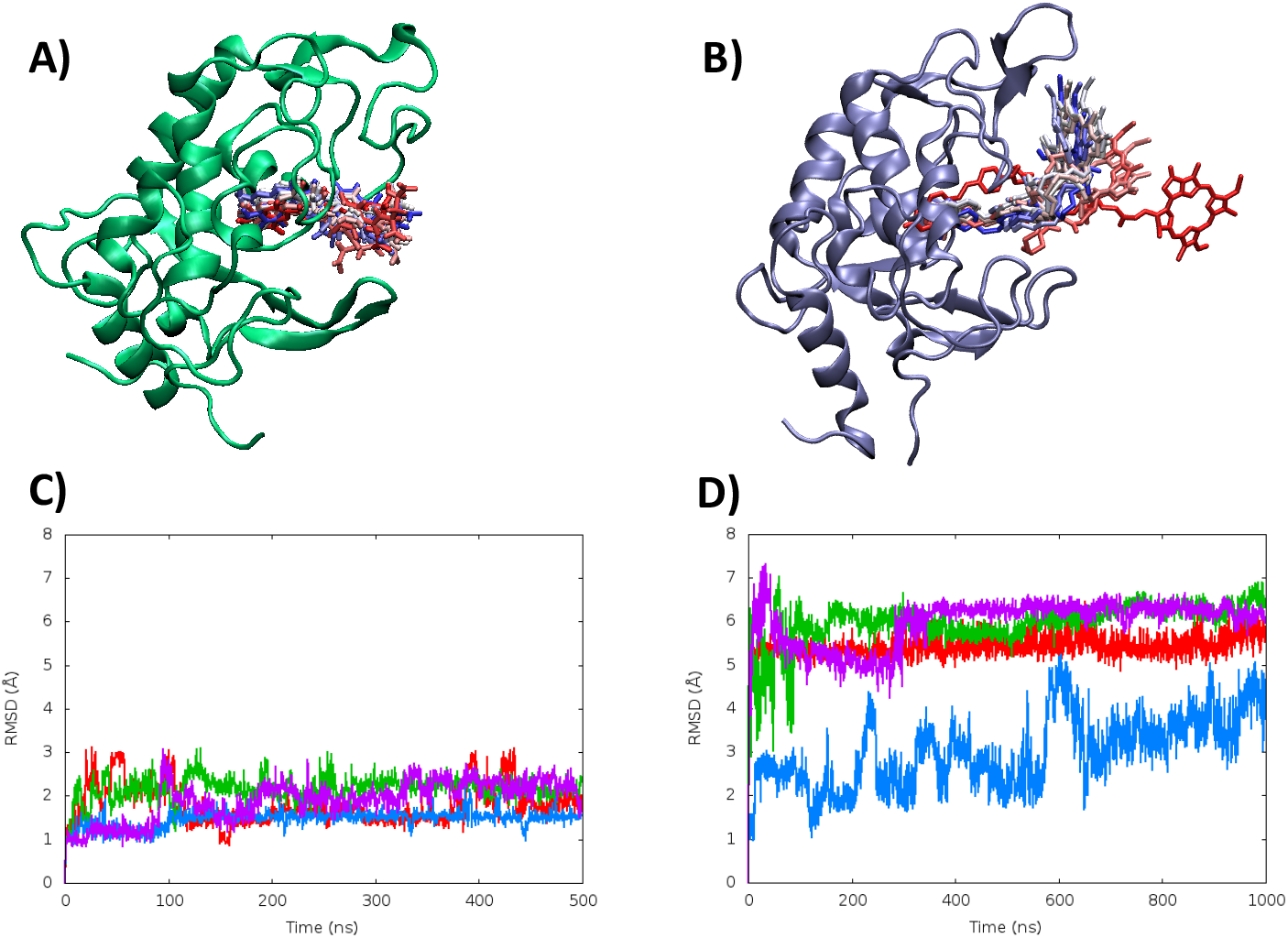
Representative structures of the FR-FA (A) and of the FR-Pyro-PEG-FA complexes (B). Note that the conformational ensemble of the ligand over the time span of the MD simulation is provided through a superposition of different structures. The corresponding time evolution of the ligand RMSD for the four replicas are reported in panel C and D, respectively.

Both adducts, i.e. the native FA and the modified FA (photosensitizer), appear stable and do not significantly deform the receptor. This can be particularly appreciated by the analysis of the RMSD for the ligands in the two systems (Figure 2 C, D) and the analysis of the time evolution of the distance between the center of mass of the protein and the ligands (Figure S1 in SI). Notably we may observe that the FA RMSD reaches a plateau at around 2 Å, while Piro-PEG-FA experiences larger deviations stabilizing at around 6 Å. Importantly; The RMSD for the protein are also reported in SI (Figure S1) confirming that the protein is stable in both conformations. Indeed, the time evolution of the FR RMSD reaches a plateau at 2 and 4 Å for the FA and Pyro-PEG-FA complex, respectively. However, while analysing the conformational ensemble explored by the ligand during the MD simulations, subtle, yet relevant differences can be observed. Indeed, Pyro-PEG-FA presents a much higher variability than FA (Figure 1). This is particularly due to conformational flexibility of the PEG linker, which can assume either straight or bent conformations. Interestingly, the flexibility of Pyro-PEG-FA appears even higher when bound to the receptor than in water, where it mostly adopts a highly linear conformation as shown in SI (Figure S19). This fact is probably due to a subtle equilibrium between the hydrophilic nature of PEG, which develops favourable interactions with the solvent water hence enforcing extended structures, and hydrophobicity of Pyro and FA, which, as it will be shown in the following sections, may develop further interactions in presence of FR, favouring bent conformations.

As also shown in SI (Figure S2) the flexibility of the protein residue, probed by the analysis of their root mean square fluctuation (RMSF) are globally conserved when FR is bound to FA or Pyro-PEG-FA with the partial exception of residue GLY143 and VAL149, which shows an increase of flexibility. However, both the structure of the binding pocket and of the whole protein are not altered by the inclusion of the photosensitizer.

In Figure 3A we report a representative structure, obtained by clustering of the MD simulation showing the stable inclusion of FA in the binding pocket. As shown in Figure 3B we may see that the ligand binding largely involves hydrogen bond interactions with ASP81 and SER101. Interestingly, further more labile electrostatic and hydrophobic interactions involving notably HIS135 and the backbone of GLY137 and TRP138 also contribute to the global stabilization of the aggregate as shown in SI (Figures S2-S10 and Table S1), even if they are not uniformly present in all the simulations. Importantly, the interaction with ASP81 and SER101, which are instead conserved in all the replicas, lock the two extremities of the ligand contributing to its affinity and selectivity for FR.

**Figure 3.**
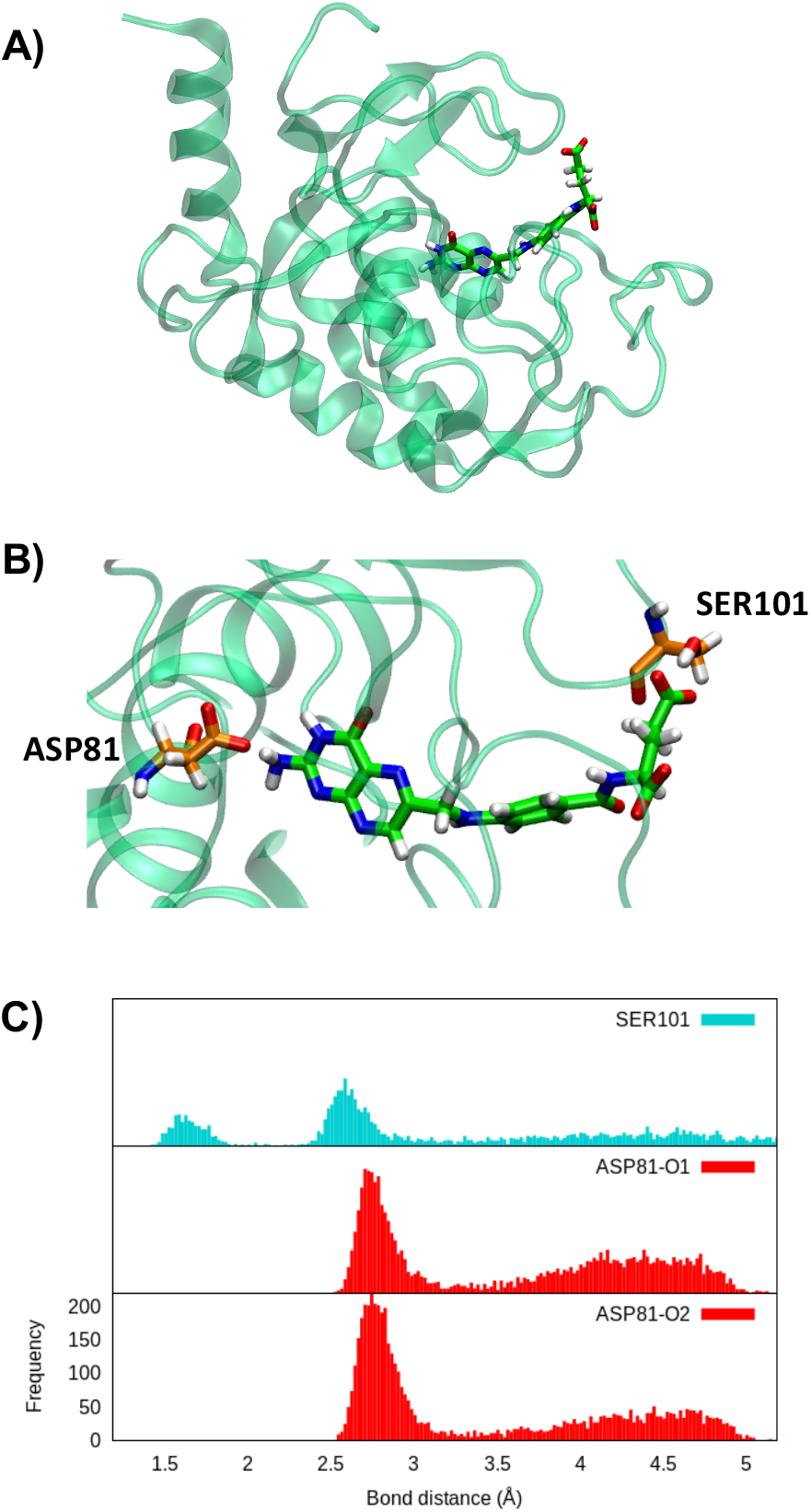
A) Representative snapshot of FA bound to FR as obtained from the MD simulation. B) Residue-scale interactions within the ligand and the protein. C) Distribution of the distances describing the interactions involving ASP81and FA, as well as SER101 and FA.

As can be seen from Figure 4, while the FA of Pyro-PEG-FA still stably and persistently interacts with ASP81 at the bottom of the binding pocket, SER101 is not engaged in significant interactions anymore. This is however partially compensated by the formation of a new interaction involving ARG103 and the amide group of FA. The absence of the locking provided by SER101 is translated in a strong mobility of the FA unit, which explores a rather large conformational space. Furthermore, a partial sliding outwards and rotation of FA inside the binding pocket can also be observed in the case of the Pyro-PEG-FA ligand. However, this movement is not sufficient to completely disrupt the main interaction with ASP81 and dissociate the complex. In addition to the main interactions common to all the replicas additional temporary electrostatic and hydrophobic interactions also develop, involving mainly ARG61 (Figure S11-S18 and Table S2)

**Figure 4.**
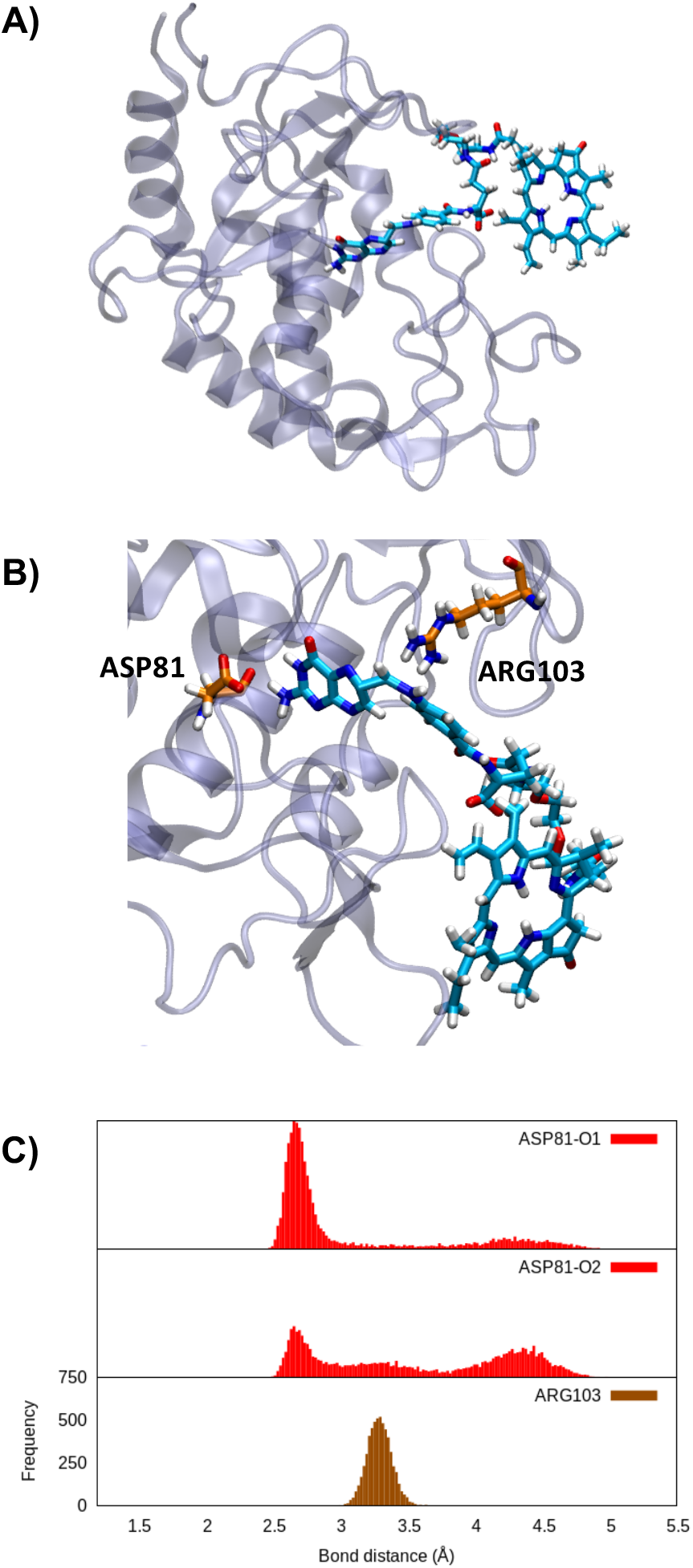
A) Representative snapshot of Pyro-PEG-FA bound to FR as obtained from the MD simulation. B) Residue-scale interactions within the ligand and the protein. C) describing the interactions involving ASP81and FA, as well as SER101 and Pyro-PEG-FA.

Since the inclusion of the Pyro-PEG tether to the FA moiety slightly changes the interaction patterns, it is reasonable to question whether the functionalized PS drug could still favourably bind to FR, and thus selectively accumulate at the surface of cancer cells. To estimate the binding affinity, we have calculated the enthalpic contribution to the binding free energies using MMPBSA and MMGBSA approaches over the MD trajectories. The calculated binding enthalpies (Figure 5) shows that the formation of the FA and Pyro-PEG-FA complexes with FR is always favourable. However, a slight difference in the binding energy can be observed: the native ligand shows a binding enthalpy estimated at about −40 (±10) kcal/mol for MMGBSA (MMPBSA) while the decorated photosensitizer is less stably bound with only −30 (±7) kcal/mol depending on the level of theory. Even if some variability can be observed between the replicas these results correlate well with the analysis of the MD trajectories, especially considering the loss of one strong interaction between FR and Pyro-PEG-FA and the larger conformational space explored by the ligand, with a partial alteration of the optimal positioning in the binding pocket. Yet, they also confirm that the proposed PS is able to selectively target FR, thus allowing the accumulation of the drug at the surface of the cancer cells.

**Figure 5.**
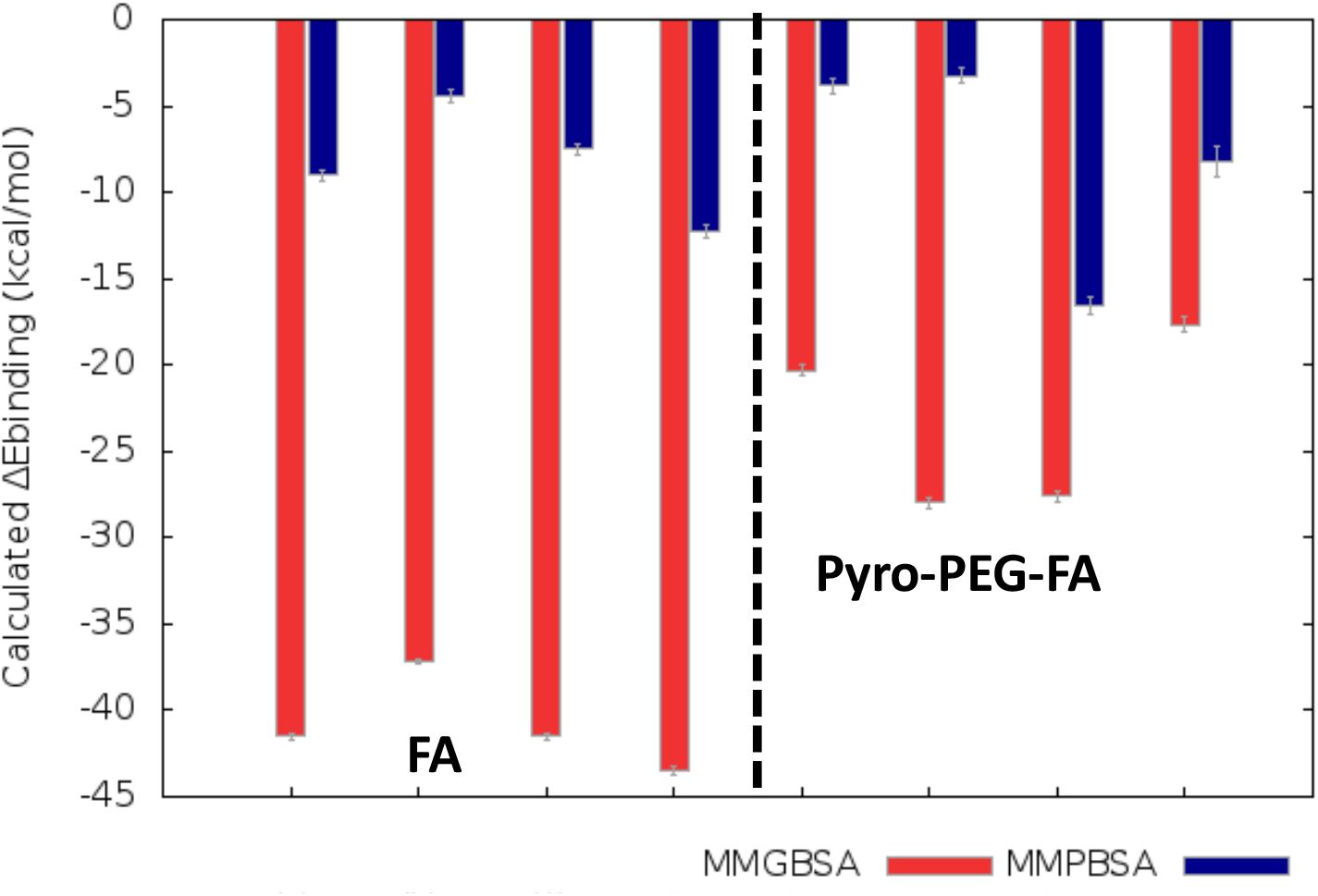
Enthalpic contribution to the binding free energy calculated for each replica of the MD simulations using MMGBSA or MMPBSA approach for the complex between FR and FA or Pyro-PEG-FA, respectively.

Finally, we have simulated the absorption spectrum of the Pyro chromophore in the Pyro-PEG-FA ligand in bulk water and in interaction with the FR to confirm that the binding with the receptor is not altering the main optical properties of the PS. The results reported in Figure 6 confirms that the spectra obtained from the two environments are almost perfectly overlapping, both in terms of bands position and shape. This is particularly true in the case of the Q-band peaking at around 500 nm, which is imperative for the potential PDT application. The analysis of the NTOs also reported in Figure 6 shows, as expected, that the transition involves the π-conjugated system of the Pyro unit. Therefore, the photophysical property of the porphyrin core, and particularly its capability to undergo intersystem crossing, should not be altered by the interaction with FR. This aspect is crucial to assure that the functionalized PS could still show an efficient population of its triplet manifolds, and thus, maintain the capacity of activate singlet oxygen.

**Figure 6.**
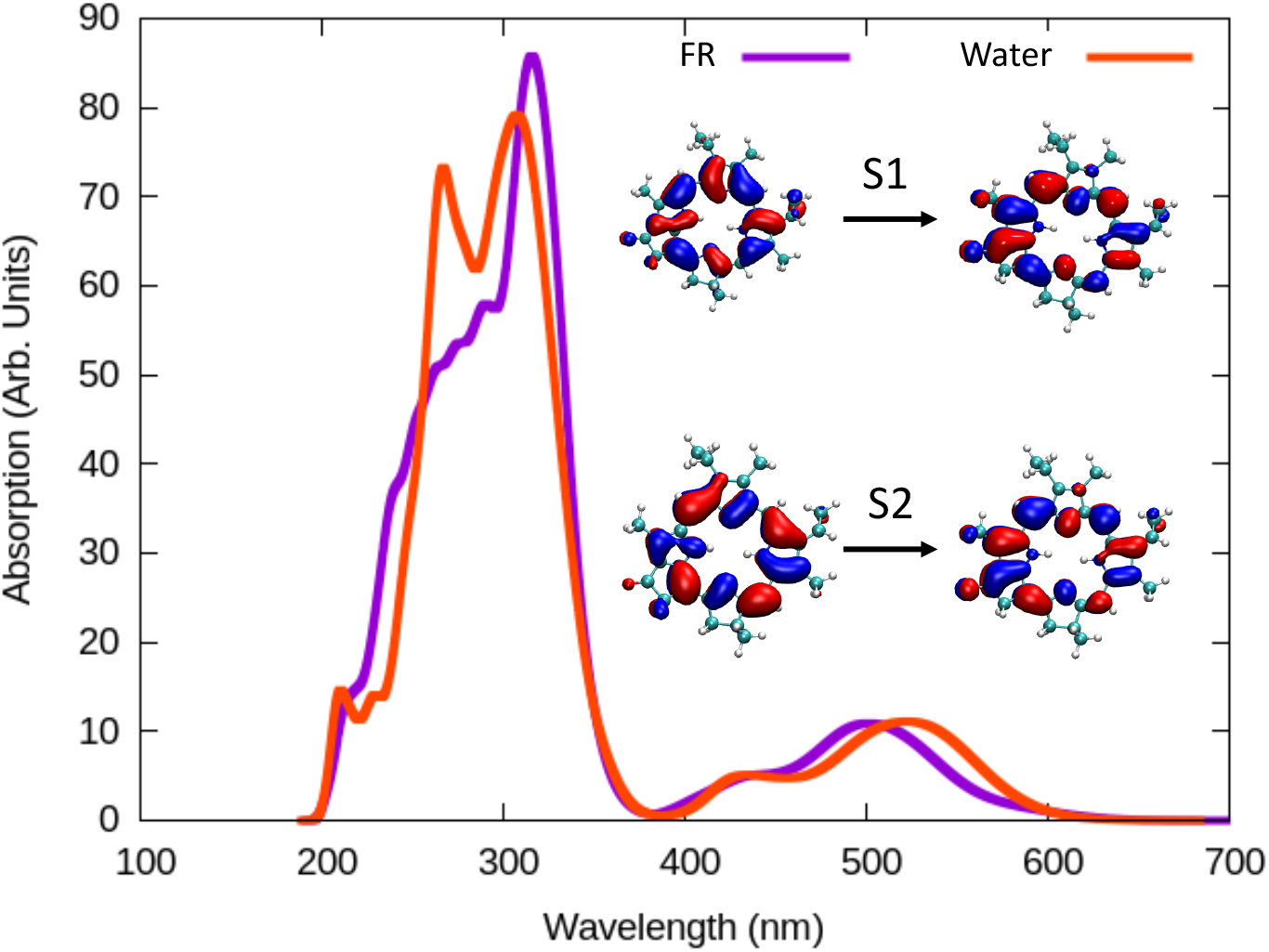
Simulated absorption spectrum of Pyro-PEG-FA in water and in interaction with FR. The NTOs for the first two excited states, encompassing the Q-band, are also given as an inlay.

## CONCLUSIONS

We have performed μ-second scale MD simulations of a functionalized PDT sensitizer (Pyro-PEG-FA), including a FA unit tethered by a PEG linker to a Pyro porphyrin unit, which can selectively interact with the human FR-α receptor. We have shown that the modified sensitizer is able to form a stable and persistent complex with the receptor through its FA-mimicking unit. Furthermore, the complex is leaving the chromophore unit (Pyro) of the PS outside of the protein core, where it can easily collide with molecular oxygen, which can thus be activated for PDT purposes. Interestingly, the protein bound PS explores a rather extended conformational space, especially thanks to the flexibility of the PEG linker. While the complex appears stable over the course of the MD simulations and the protein structure and flexibility profile are not significantly perturbed, we have shown that the interaction patterns change and are slightly reduced compared to the native ligand. This in turn translates to an increased mobility of the Pyro-PEG-FA ligand and a reduced binding free energy. However, the global stability of the complex clearly points toward the possible incorporation of the PS to the receptor, which should result in its accumulation at the surface of FR-expressing cells. Finally, we have shown that the optical properties of the FR-bound sensitizer are not altered compared to those expressed in bulk water, suggesting that the intersystem crossing capability and ultimately singlet oxygen quantum yield should not be affected. Therefore, we have confirmed that the linkage of a PS with a known ligand targeting a receptor, which is overexpressed in cancer cell may be utilized as a most valuable strategy for active drug delivery in PDT allowing the selective accumulation of the sensitizer to the surface of cancer cells. This approach can be particularly attractive for the use of PDT as an adjuvant strategy during cancer surgery. Considering that FR is highly overexpressed in case of ovarian cancer, the present PDT strategy can be particularly exploited to treat potential abdominal metastasis via laparoscopic approaches [76].

Future studies that continue the exploration of active PDT delivery, in particular considering a membrane bound FR receptor to assess the proximity of the PS to the cellular membrane, which could represent the singlet oxygen target are currently underway. We also plan to propose rational design of the linker to optimize the binding affinity of the PS with the receptor and to consider other receptors to selectively target other cancer cells. This will be particularly attractive in the case of pancreatic adenocarcinoma for which conventional therapeutic approaches are mostly inefficient.

## ACKNOWLEDGEMENTS

The authors thank GENCI, Explor and National Center for High Performance Computing of Turkey (UHeM) under grant number 1011062021, computing centers and the Platform P3MB for computational resources. The authors thanks ANR and CGI for their financial support of this work through Labex SEAM ANR 11 LABEX 086, ANR 11 IDEX 05 02 and PIRATE. The support of the IdEx “Université Paris 2019” ANR-18-IDEX-0001. Support from the PEPR LUMA is also gratefully acknowledged. BKF and SC thank TUBITAK (Project Number: 120Z659) for financial support. BKF would also like to thank the French Embassy in Turkey for a joint PhD grant, as well as TUBITAK for the 2214A International Research Fellowship Programme for PhD Students.

