## Supplementary Information for "In Silico Study of Active Delivery of a Photodynamic Therapy Drug Targeting the Folate Receptor"

### LIST OF FIGURES


FIGURE S1 ROOT-MEAN-SQUARE-DEVIATION ANALYSIS OF THE PROTEIN BACKBONE DURING THE MD SIMULATIONS IN THE PRESENCE OF FA (LEFT) AND PYRO-PEG-FA (RIGHT)..........S3
FIGURE S2. ROOT-MEAN-SQUARE-FLUCTUATION ANALYSIS THROUGHOUT THE TRAJECTORIES OF FIRST REPLICA OF SIMULATIONS. ..........S3
FIGURE S3. DISTRIBUTION OF THE DISTANCES OF MAIN INTERACTIONS BETWEEN FOLATE RECEPTOR AND FOLATE OCCURRED IN REPLICA 1..........S5
FIGURE S4. DISTRIBUTION OF THE DISTANCES OF MAIN INTERACTIONS BETWEEN FOLATE RECEPTOR AND FOLATE OCCURRED IN REPLICA 2..........S7
FIGURE S5. DISTRIBUTION OF THE DISTANCES OF MAIN INTERACTIONS BETWEEN FOLATE RECEPTOR AND FOLATE OCCURRED IN REPLICA 3..........S9
FIGURE S6. DISTRIBUTION OF THE DISTANCES OF MAIN INTERACTIONS BETWEEN FOLATE RECEPTOR AND FOLATE OCCURRED IN REPLICA 4..........S11
FIGURE S7. DISTRIBUTION OF THE DISTANCES OF MAIN INTERACTIONS BETWEEN FOLATE RECEPTOR AND PYRO-PEG-FA OCCURRED IN REPLICA 1. ..........S14
FIGURE S8. DISTRIBUTION OF THE DISTANCES OF MAIN INTERACTIONS BETWEEN FOLATE RECEPTOR AND PYRO-PEG-FA OCCURRED IN REPLICA 2. ..........S16
FIGURE S9. DISTRIBUTION OF THE DISTANCES OF MAIN INTERACTIONS BETWEEN FOLATE RECEPTOR AND PYRO-PEG-FA OCCURRED IN REPLICA 3. ..........S17
FIGURE S10. DISTRIBUTION OF THE DISTANCES OF MAIN INTERACTIONS BETWEEN FOLATE RECEPTOR AND PYRO-PEG-FA OCCURRED IN REPLICA 4. ..........S19
FIGURE S11. ROOT-MEAN-SQUARE-DEVIATION OF PYRO-PEG-FA IN BULK WATER, AND ITS REPRESENTATIVE STRUCTURE. ..........S20
FIGURE S12. COMPARATIVE TIME EVALUATION OF COM DISTANCE OF FOLATE AND PYRO UNITS WHEN INTERACTING WITH THE FOLATE RECEPTOR (RED) AND IN BULK WATER (BLUE). ...S20


### LIST OF TABLES


TABLE S1. PERSISTENT INTERACTIONS BETWEEN FOLATE RECEPTOR AND FOLATE. ..........S4
TABLE S2. TIME EVALUATION OF MAIN INTERACTIONS BETWEEN FOLATE RECEPTOR AND FOLATE OCCURRED IN REPLICA 1..........S6
TABLE S3. TIME EVALUATION OF MAIN INTERACTIONS BETWEEN FOLATE RECEPTOR AND FOLATE OCCURRED IN REPLICA 2..........S8
TABLE S4. TIME EVALUATION OF MAIN INTERACTIONS BETWEEN FOLATE RECEPTOR AND FOLATE OCCURRED IN REPLICA 3..........S10
TABLE S5. TIME EVALUATION OF MAIN INTERACTIONS BETWEEN FOLATE RECEPTOR AND FOLATE OCCURRED IN REPLICA 4..........S12
TABLE S6. PERSISTENT INTERACTIONS BETWEEN FOLATE RECEPTOR AND PHOTOSENSITIZER. S13
TABLE S7. TIME EVALUATION OF MAIN INTERACTIONS BETWEEN FOLATE RECEPTOR AND PYRO-PEG-FA OCCURRED IN REPLICA 1. ..........S15
TABLE S8. TIME EVALUATION OF MAIN INTERACTIONS BETWEEN FOLATE RECEPTOR AND PYRO-PEG-FA OCCURRED IN REPLICA 2. ..........S16
TABLE S9. TIME EVALUATION OF MAIN INTERACTIONS BETWEEN FOLATE RECEPTOR AND PYRO-PEG-FA OCCURRED IN REPLICA 3. ..........S18
TABLE S10. TIME EVALUATION OF MAIN INTERACTIONS BETWEEN FOLATE RECEPTOR AND PYRO-PEG-FA OCCURRED IN REPLICA 4. ..........S19

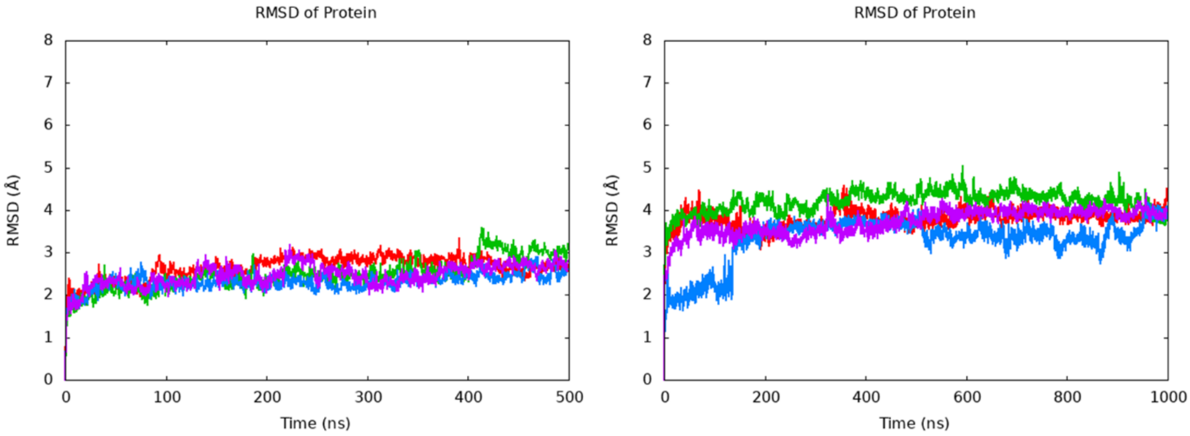


Figure S1 Root-Mean-Square-Deviation Analysis of the protein backbone during the MD simulations in the presence of FA (left) and Pyro-PEG-FA (right).

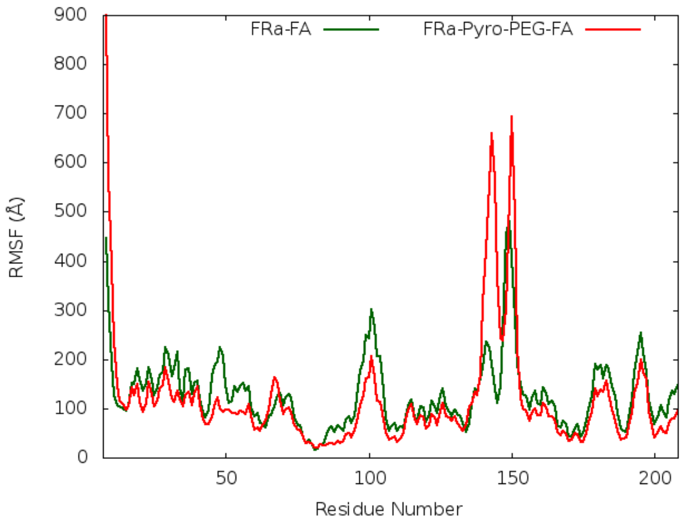


Figure S2. Root-Mean-Square-Fluctuation analysis throughout the trajectories of first replica of simulations.

Table S1. Persistent interactions between Folate receptor and Folate.

| FRα | FA | REP1 | REP2 | REP3 | REP4 |
|---|---|---|---|---|---|
| TRP140@NE1-HE1 | @O5 | 63% | n/a | 63% | 12% |
| SER101@OG-HG | @O4 | 45% | 18% | 45% | 15% |
| ASP81@OD1 | @N2-H7 | 37% | 17% | 48% | 63% |
| ASP81@OD2 | @N2-H7 | 22% | 18% | n/a | n/a |
| ASP81@OD1 | @N2-H16 | 21% | 45% | n/a | 18% |
| GLY137@N-H | @O6 | 36% | n/a | 45% | n/a |
| HIE135@NE2-HE2 | @O1 | 30% | n/a | n/a | 21% |
| TRP138@N-H | O5 | 25% | n/a | 27% | n/a |
| THR82@OG1 | @N2-H16 | 12% | 10% | 22% | 12% |
| ARG103@NH1-HH11 | @O1 | n/a | n/a | 19% | n/a |
| ARG61@NE-HE | @O6 | n/a | 25% | n/a | n/a |

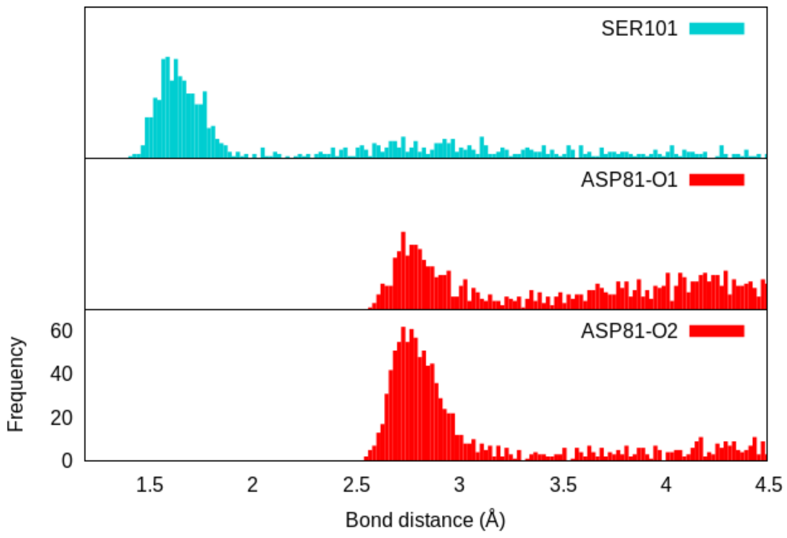


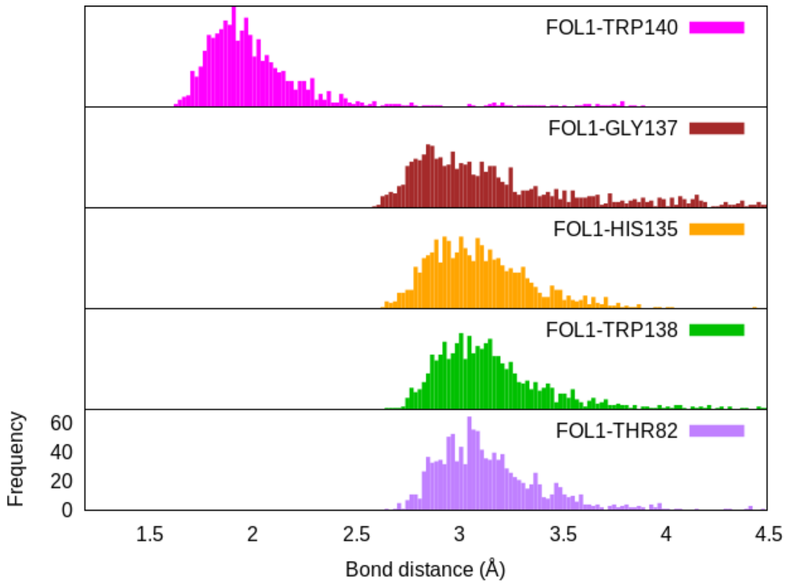


Figure S3. Distribution of the distances of main interactions between folate receptor and folate occurred in Replica 1

Table S2. Time evaluation of main interactions between folate receptor and folate occurred in Replica 1.

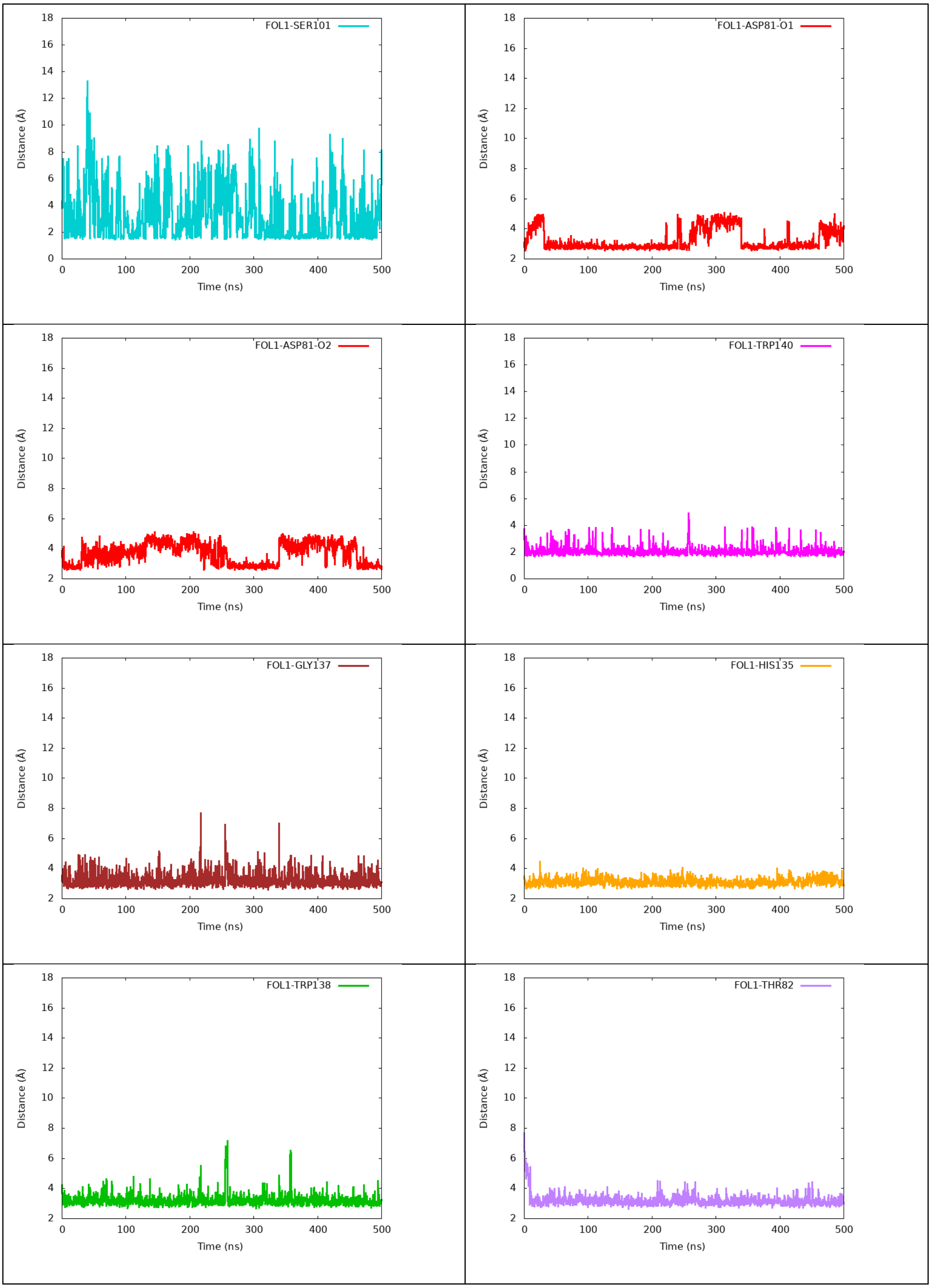

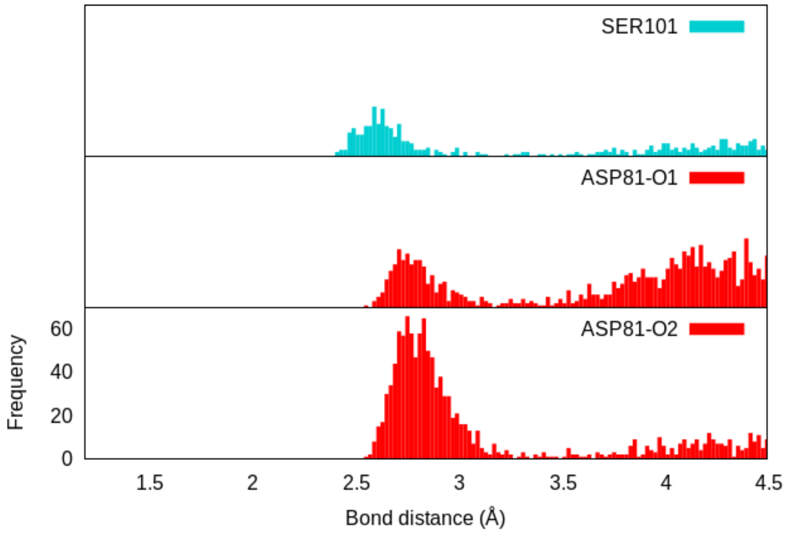


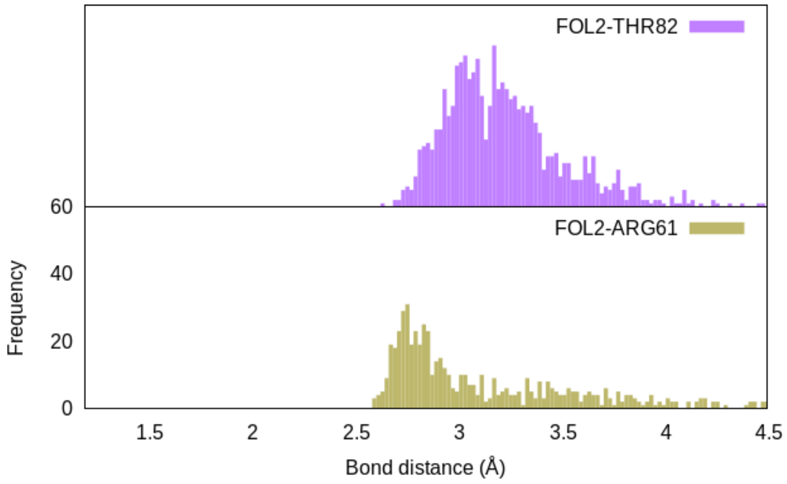


Figure S4. Distribution of the distances of main interactions between folate receptor and folate occurred in Replica 2.

Table S3. Time evaluation of main interactions between folate receptor and folate occurred in Replica 2.

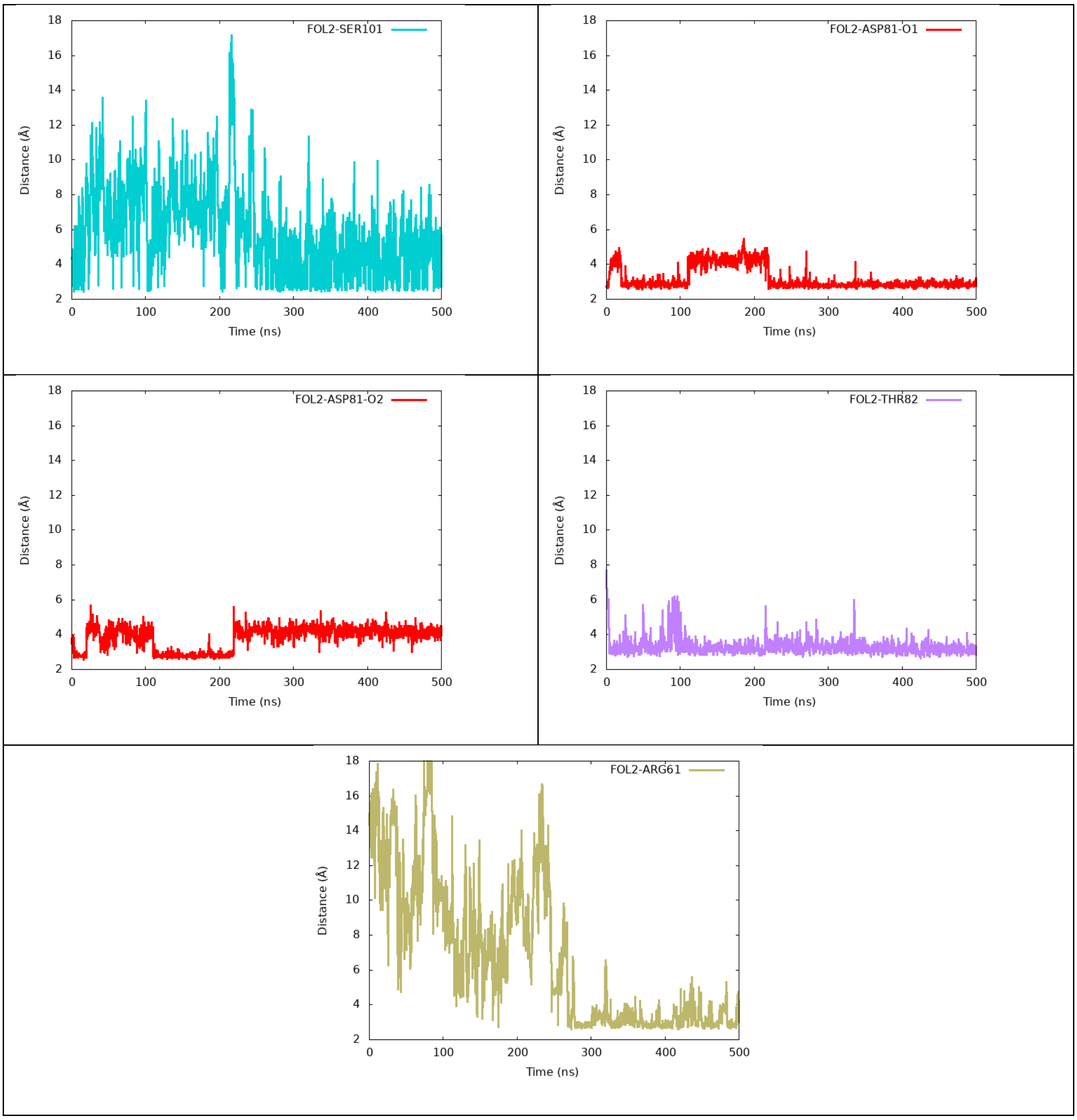

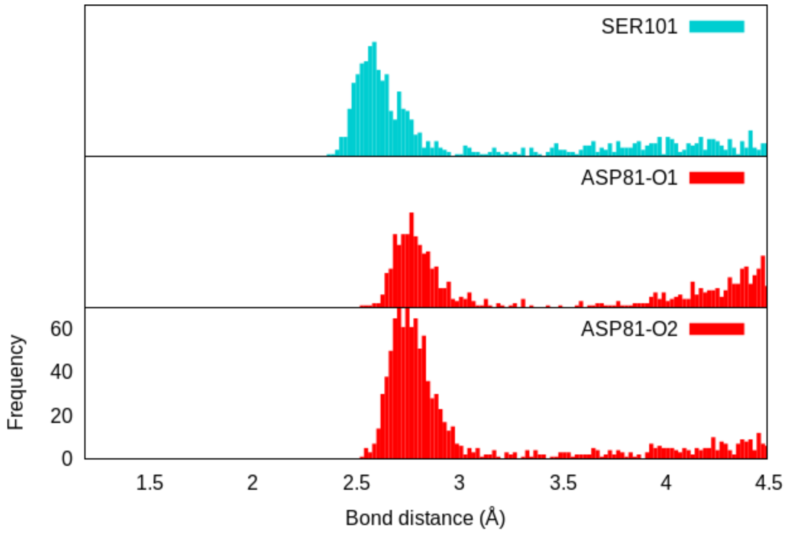


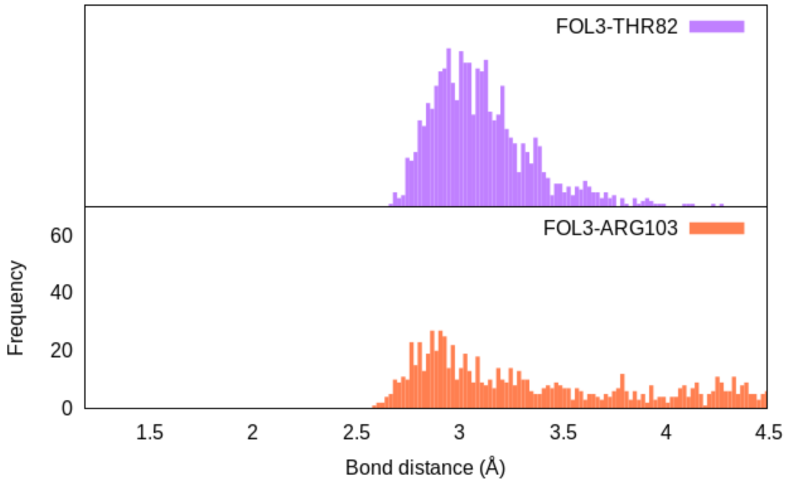


Figure S5. Distribution of the distances of main interactions between folate receptor and folate occurred in Replica 3.

Table S4. Time evaluation of main interactions between folate receptor and folate occurred in Replica 3.

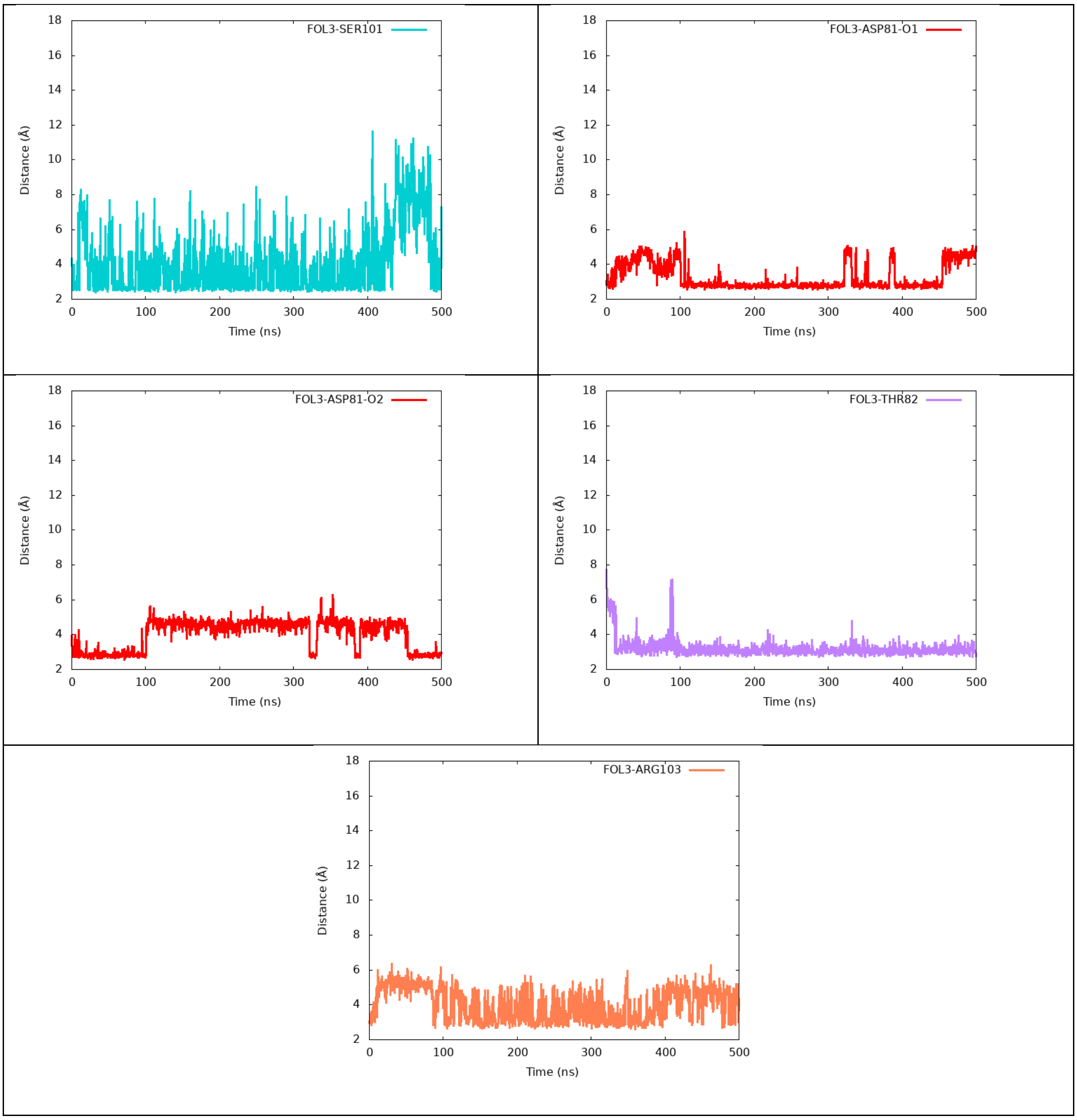

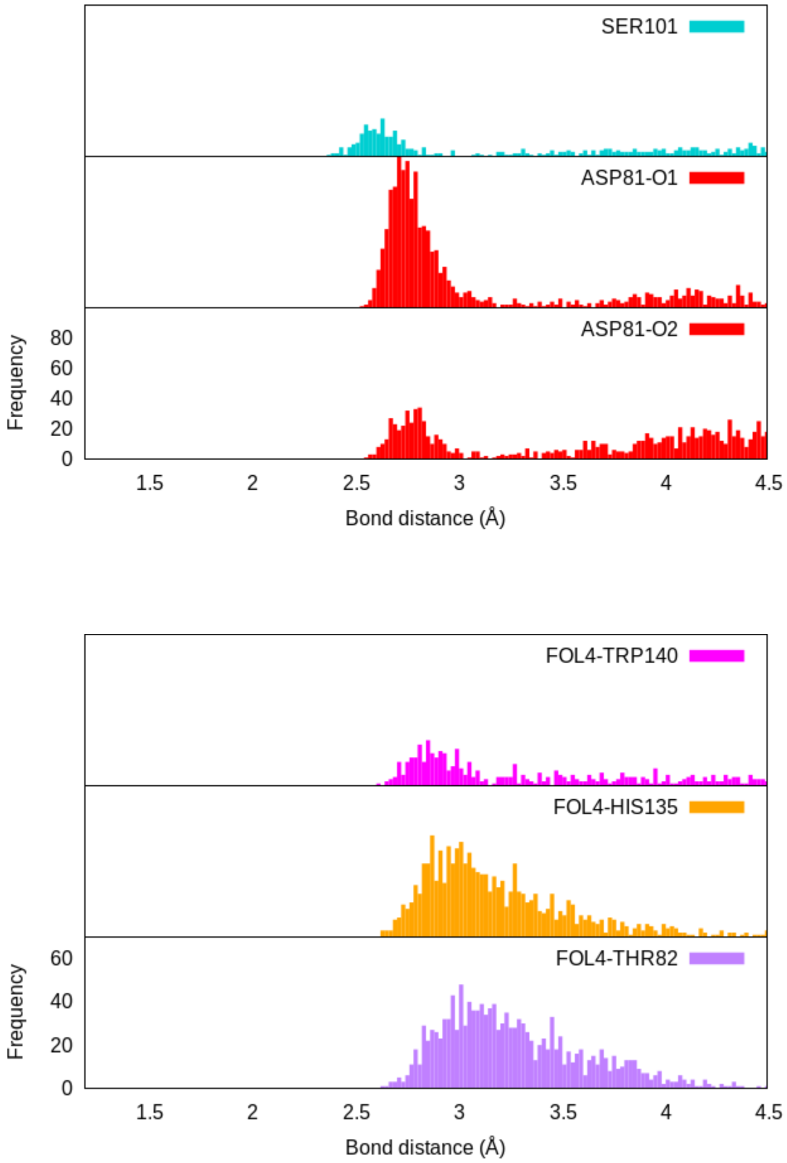


Figure S6. Distribution of the distances of main interactions between folate receptor and folate occurred in Replica 4.

Table S5. Time evaluation of main interactions between folate receptor and folate occurred in Replica 4.

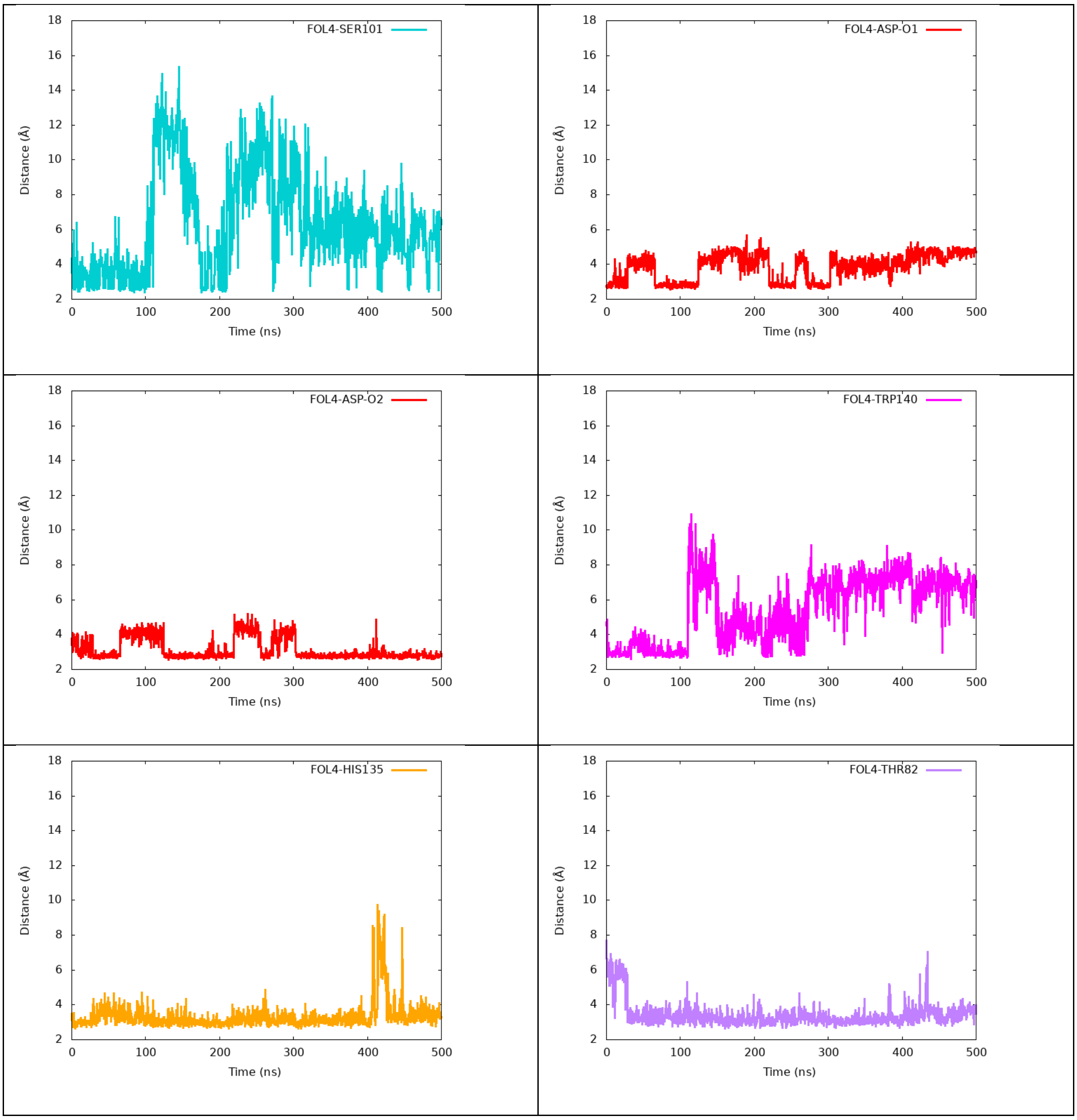

Table S6. Persistent interactions between Folate receptor and photosensitizer.

| **FRα** | **Pyro-PEG-FA** | **REP1** | **REP2** | **REP3** | **REP4** |
|---|---|---|---|---|---|
| ARG103@NH1-HH11 | @N6 | 89% | n/a | 78% | n/a |
| ARG103@NH2-HH21 | @N6 | 84% | n/a | 94% | n/a |
| ARG61@NE-HE | @O3 | 56% | n/a | n/a | n/a |
| ASP81@OD2 | @N2-H7 | 35% | n/a | 21% | 20% |
| ASP81@OD1 | @N2-H7 | 20% | n/a | 71% | 85% |
| ASP81@O | @N2-H7 | n/a | 36% | 21% | n/a |
| ASP81@OD2 | @N2-H16 | n/a | 17% | n/a | n/a |
| ASP81@OD1 | @N2-H16 | 20% | 33% | n/a | 74% |
| ASP81@O | @N2-H16 | n/a | 29% | n/a | n/a |

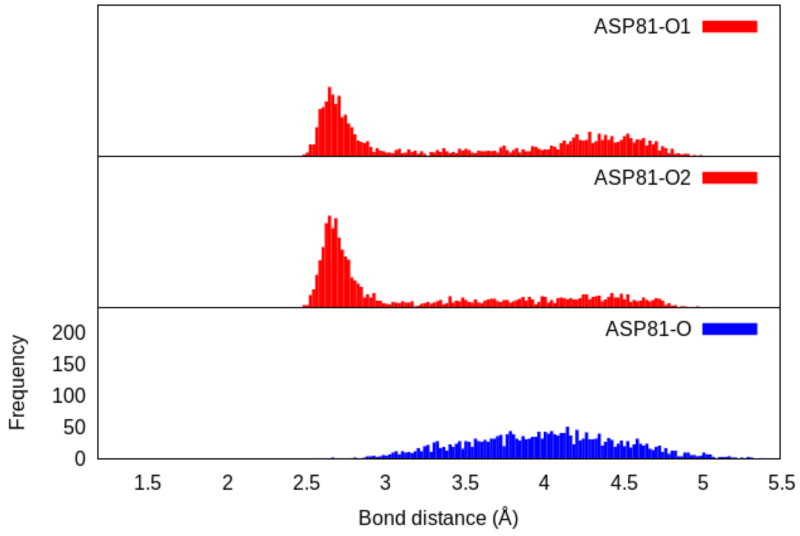


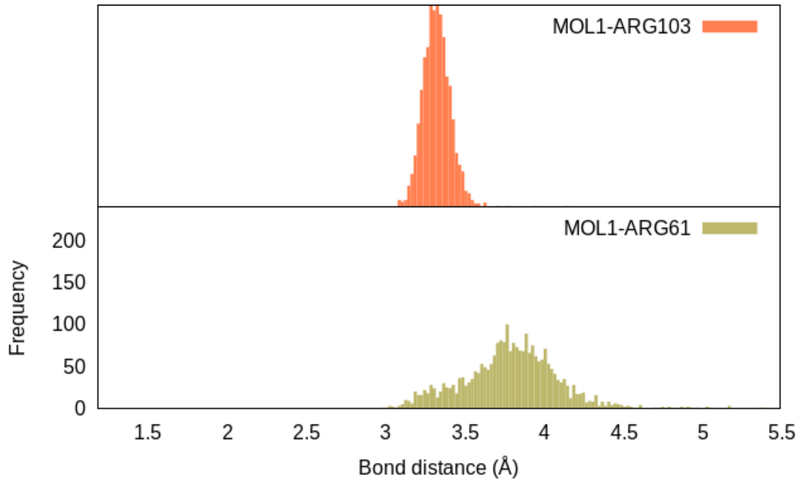


Figure S7. Distribution of the distances of main interactions between Folate Receptor and Pyro-PEG-FA occurred in Replica 1.

Table S7. Time evaluation of main interactions between Folate Receptor and Pyro-PEG-FA occurred in Replica 1.

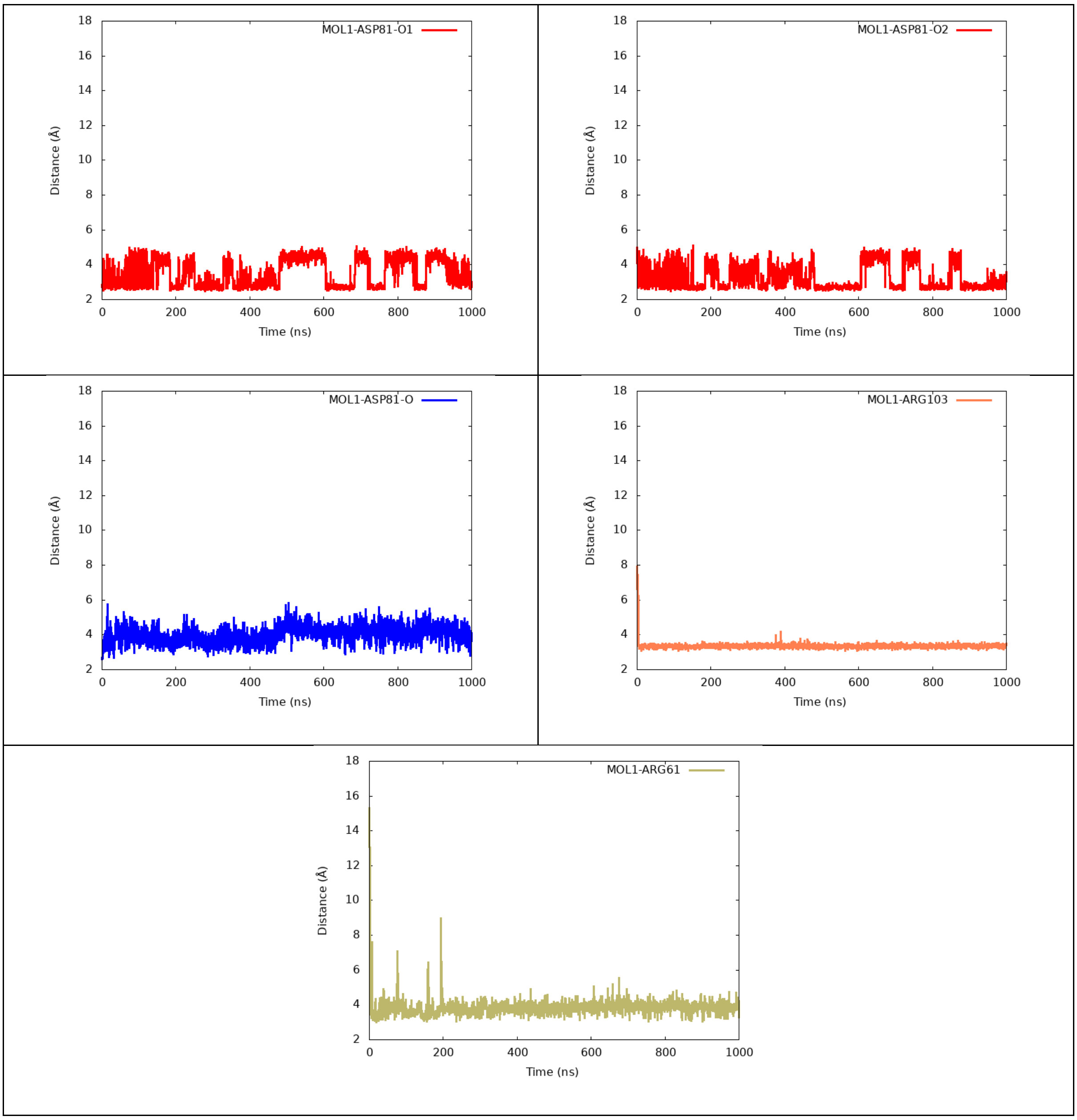

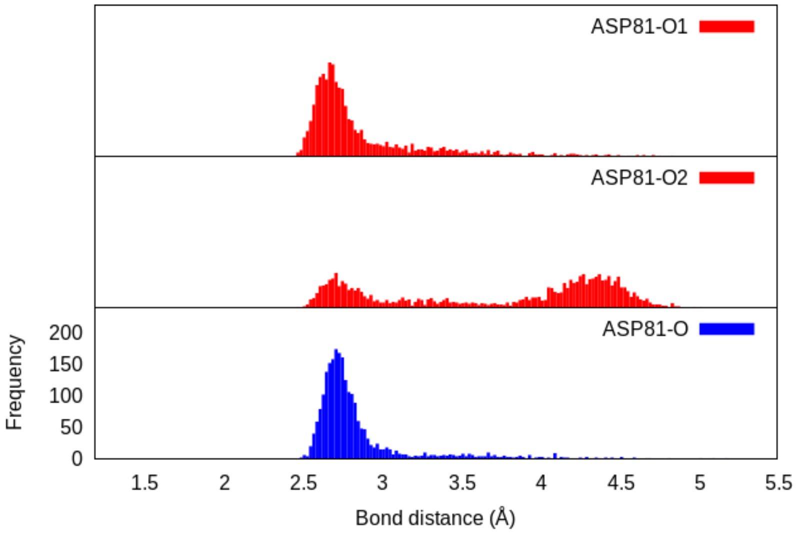


Figure S8. Distribution of the distances of main interactions between Folate Receptor and Pyro-PEG-FA occurred in Replica 2.

Table S8. Time evaluation of main interactions between Folate Receptor and Pyro-PEG-FA occurred in Replica 2.

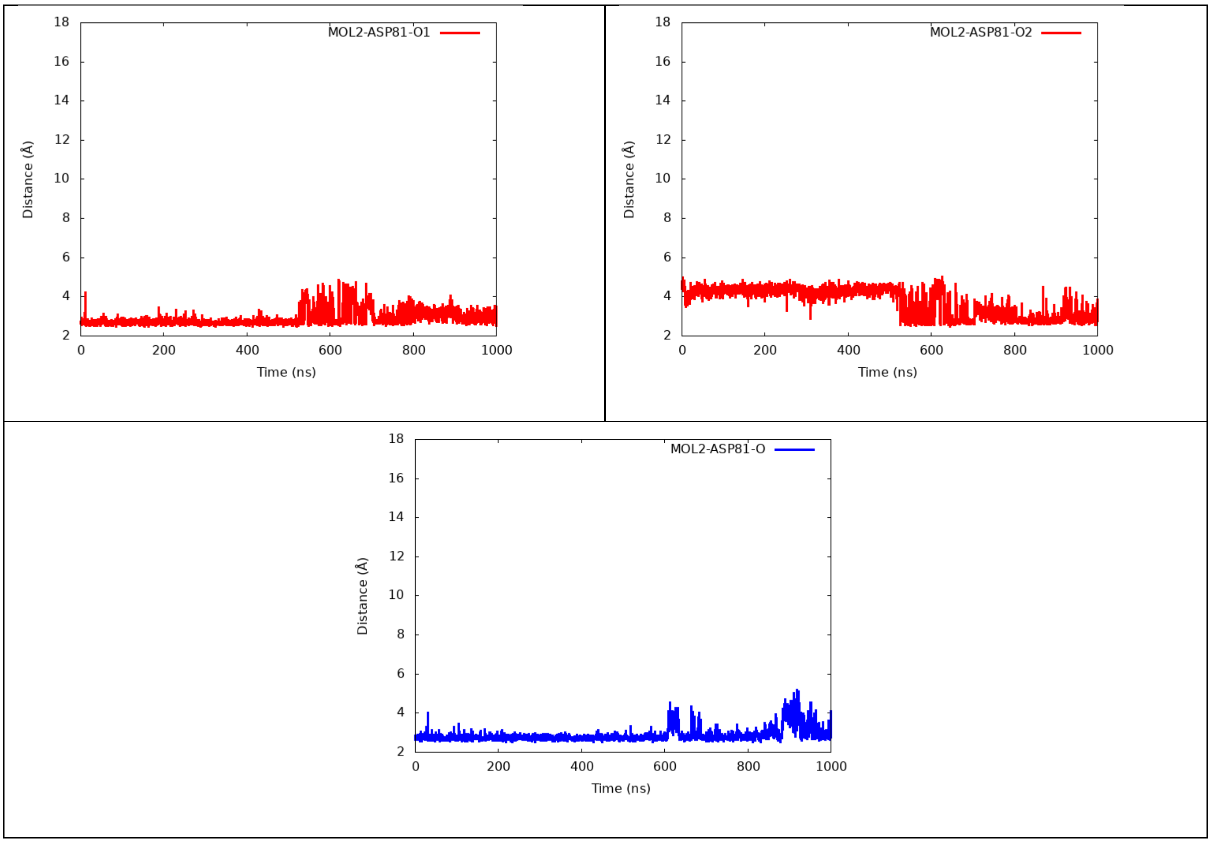

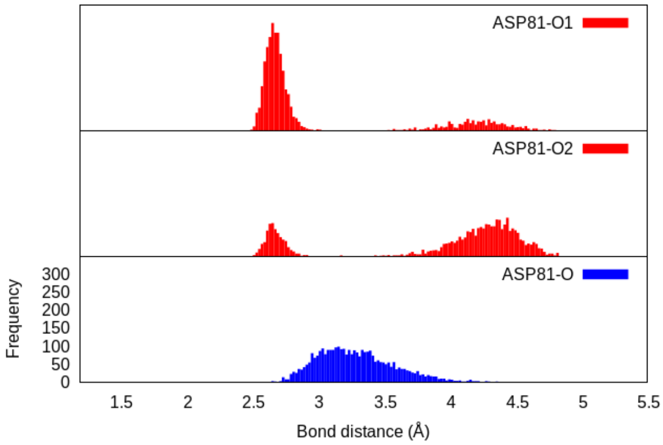


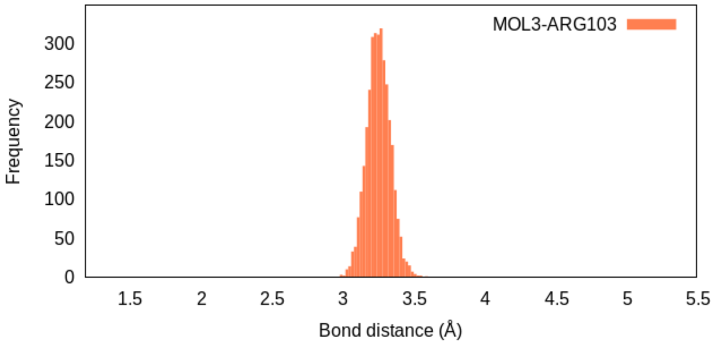


Figure S9. Distribution of the distances of main interactions between Folate Receptor and Pyro-PEG-FA occurred in Replica 3.

Table S9. Time evaluation of main interactions between Folate Receptor and Pyro-PEG-FA occurred in Replica 3.

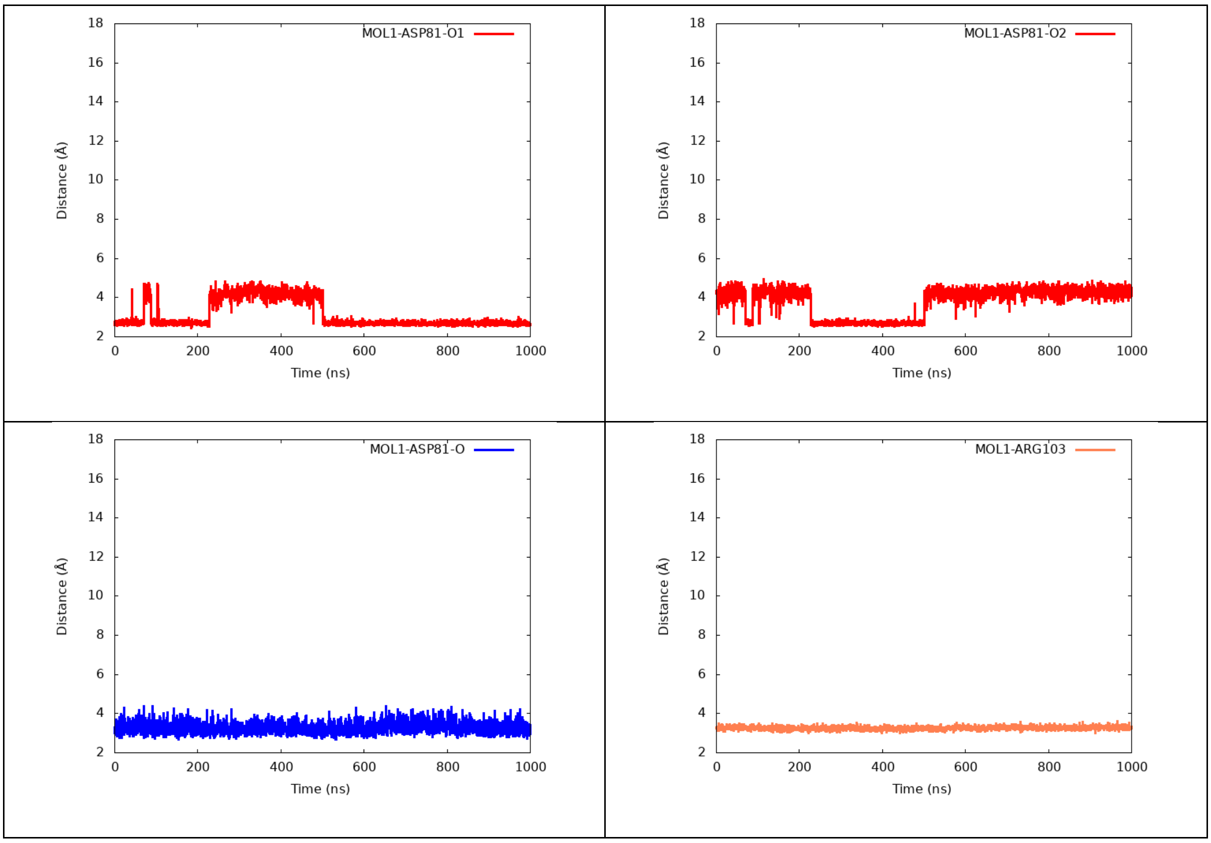

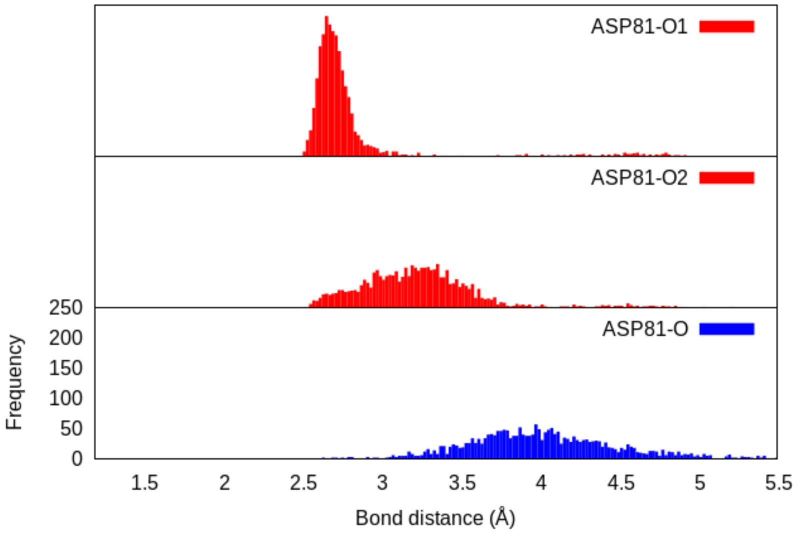


Figure S10. Distribution of the distances of main interactions between Folate Receptor and Pyro-PEG-FA occurred in Replica 4.

Table S10. Time evaluation of main interactions between Folate Receptor and Pyro-PEG-FA occurred in Replica 4.

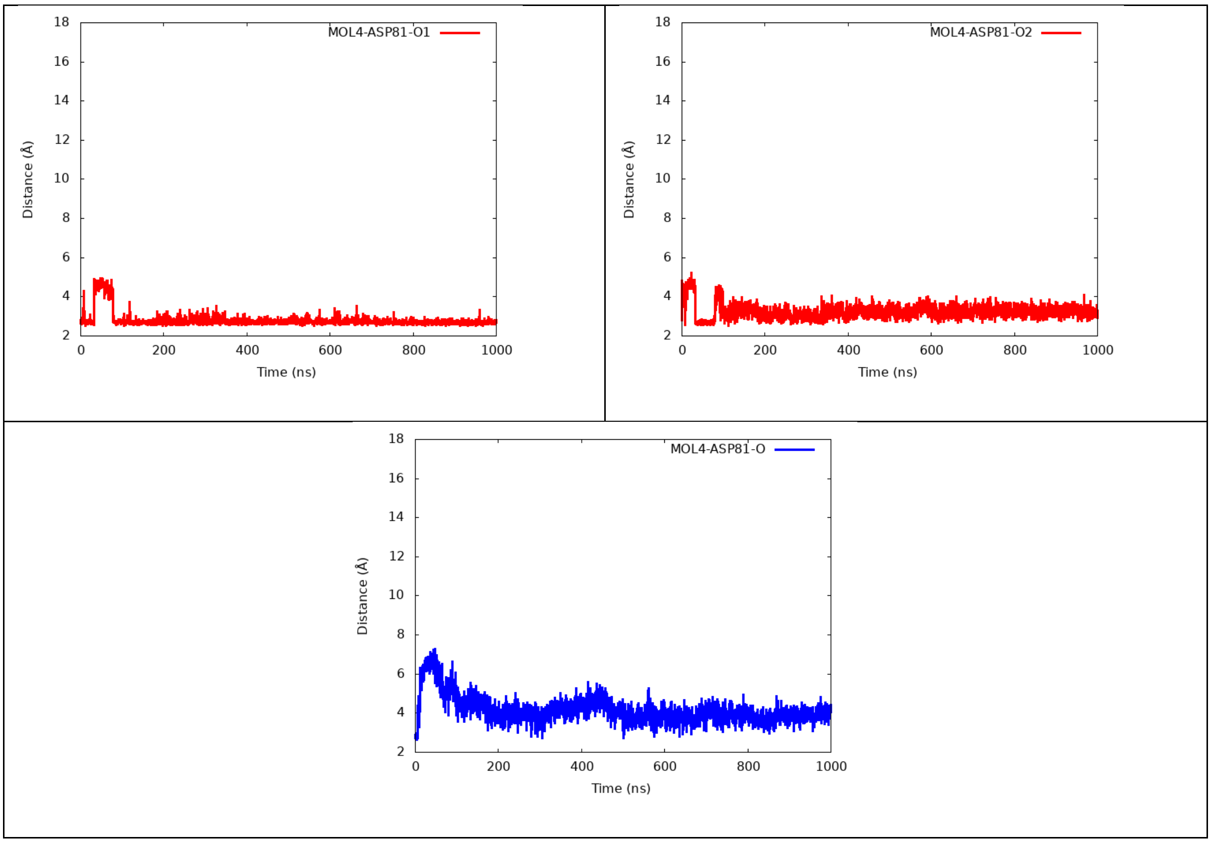

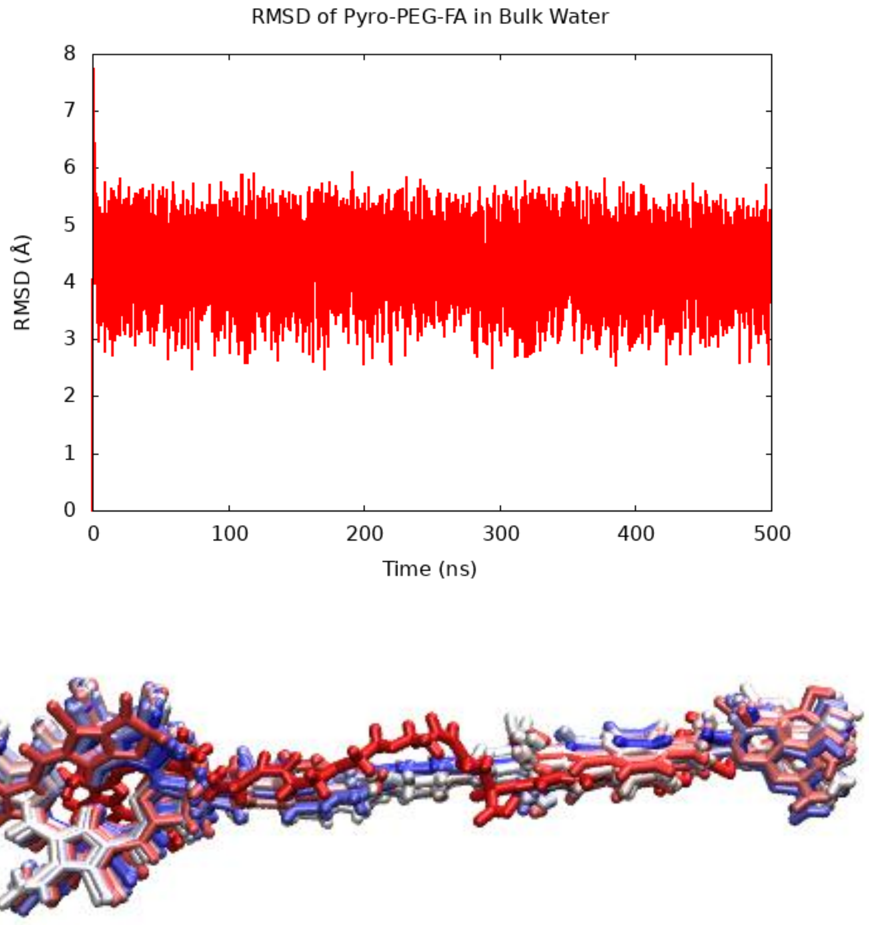


Figure S11. Root-Mean-Square-Deviation of Pyro-PEG-FA in Bulk Water, and its representative structure.

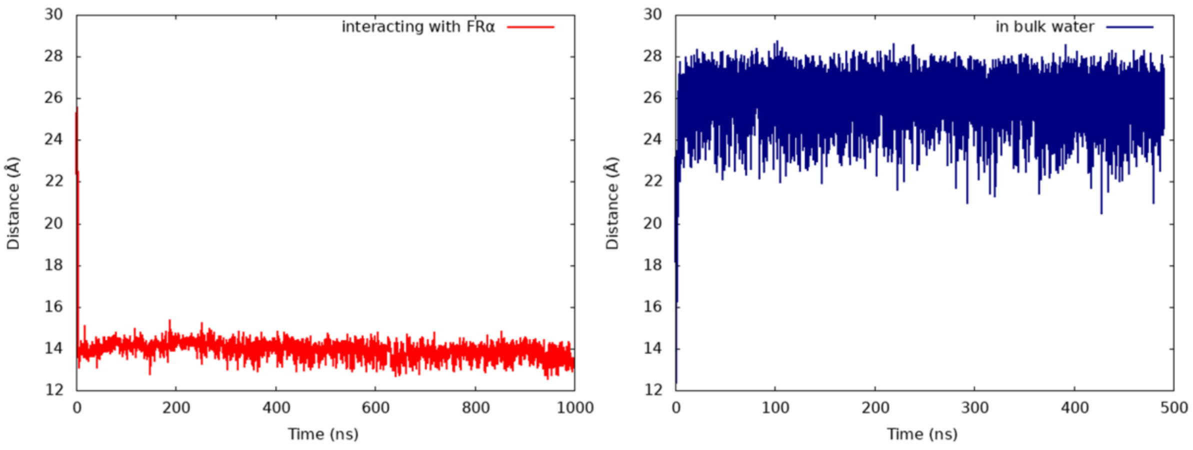


Figure S12. Comparative time evaluation of COM distance of Folate and Pyro units when interacting with the Folate Receptor (red) and in bulk water (blue).
